# Enhancing Mycobacterial Clearance via Heterologous BCG Prime Boost Strategy Using Recombinant *Mycobacterium smegmatis*

**DOI:** 10.64898/2026.08.31.748203

**Authors:** HS Ji, KR Kang, JA Kim, YH Kwon, HM Kang, UY Choi, JH Kang, GS Choi

**Author notes:** Corresponding author: (Choi GS). These authors contributed equally to this work (HS Ji, KR Kang).

## Abstract

*Bacille Calmette–Guérin* (BCG) remains the only licensed tuberculosis (TB) vaccine, yet its protective efficacy against adult pulmonary TB is limited. To overcome this, we evaluated a novel recombinant *Mycobacterium smegmatis* (rMs) booster vaccine expressing Ag85B and ESAT-6 via the pMyong2 vector, combined with oat-derived β-glucan adjuvant, to enhance BCG-primed immunity. BALB/c mice were primed with BCG and boosted twice with homologous BCG or varying doses of rMs with or without β-glucan. Five weeks post-booster, mice were challenged intraperitoneally with *M. tuberculosis* H37Ra. Systemic bacterial clearance in liver tissue, cell-mediated immunity IFN-γ, IL-17a, and antibody responses IgG, IgG2c, IgA were evaluated 3 weeks post-challenge. All rMs booster groups (rMs, high-dose rMs H, low-dose with β-glucan, rMs β) achieved complete systemic clearance of H37Ra with no detectable CFUs (colony forming unit), significantly outperforming mock-vaccinated and repeated BCG groups (\*\**p* < 0.01). Repeated BCG revaccination showed higher bacterial loads than single BCG (\*\*\**p* < 0.001), confirming no booster effect. Low-dose rMs with β-glucan maintained 100% survival and total bacterial elimination despite a 50% dose reduction. Immune markers did not strictly correlate with clearance. Homologous BCG revaccination provides no additive protection, whereas pMyong2-based rMs confers robust systemic bacterial clearance. Adjuvanting with β-glucan enables dose reduction while preserving maximum efficacy, establishing rMs as a promising heterologous TB booster candidate.

## Introduction

Tuberculosis (TB), a representative chronic respiratory infectious disease caused by *Mycobacterium tuberculosis (M. tuberculosis)* infection, remains one of the most critical public health challenges worldwide. According to the World Health Organization (WHO), an estimated 10.8 million people developed TB and approximately 1.25 million died from the disease globally in 2023 (1). In particular, the persistent emergence of multidrug-resistant TB (MDR-TB) and the widespread prevalence of latent TB infection (LTBI) serve as primary obstacles to the global eradication of tuberculosis (2).

The Bacille Calmette–Guérin (BCG) vaccine, derived from *Mycobacterium bovis*, was introduced in 1921 and remains the only currently available vaccine for the prevention of tuberculosis (TB). The World Health Organization (WHO) recommends a single dose of BCG vaccination for all healthy neonates immediately after birth in countries or settings with a high incidence of TB (3). The BCG vaccine is well established to be effective in preventing severe forms of pediatric tuberculosis, such as miliary TB and tuberculous meningitis (4), thereby playing a critical role in reducing pediatric TB mortality in endemic regions.

However, the BCG vaccine exhibits limited efficacy in preventing pulmonary tuberculosis (TB) in adults, with protective effects varying significantly across geographic regions and population groups (5). Previous studies have reported substantial variability in the efficacy of BCG against adult pulmonary TB, with notably lower protection observed in tropical and subtropical regions (6). This variability has been attributed to multiple factors, including host genetic background, nutritional and immunological status, exposure to environmental non-tuberculous mycobacteria (NTM), variations among BCG strains, and waning immunity over time following vaccination. In particular, exposure to NTMs can induce cross-immunity through shared antigens with BCG; however, under certain conditions, it has been reported to block or alter BCG-induced immune responses, thereby diminishing protective efficacy (7). To overcome these limitations, diverse next-generation TB vaccine strategies are currently being developed (8). Nevertheless, there remains a need for new vaccine strategies that can complement BCG and improve protection against tuberculosis. Although many TB vaccine candidates are being developed to boost BCG-induced immunity or enhance cell-mediated responses against TB-specific antigens, further improvements in the magnitude and durability of protective immunity are still required. Consequently, the development of novel heterologous booster vaccine platforms that can enhance BCG-primed immunity remains an important objective in next-generation TB vaccine research (9).

In this study, we evaluated non-clinically the efficacy of a recombinant vector vaccine—termed the pMyong2 vector vaccine—engineered using *Mycobacterium smegmatis* (*M. smegmatis*), a fast-growing NTM that is easy to manipulate genetically, to express the tuberculosis-specific antigens Ag85B and ESAT-6. Ag85B, a major secreted antigen belonging to the mycolyl transferase family, is a representative antigen that induces TB-specific T-cell immunity. ESAT-6 is a highly immunogenic antigen located in the RD1 region; since it is absent in BCG, it holds strong potential for use as a BCG booster antigen. By utilizing oat-derived β-glucan as an adjuvant, we evaluated the preclinical booster effect of this recombinant *M. smegmatis* vector vaccine following BCG vaccination. Through this approach, we aimed to evaluate the preclinical potential of a novel mycobacterial vector-based TB booster platform to enhance protective immunity and mycobacterial clearance following BCG priming.

## Materials and Methods

### Mycobacterium Cultivation

The *M. smegmatis* strain carrying the recombinant *M. smegmatis* vector (rMs; Fig. 1) (10) was developed, manufactured by CLIPS BnC Co., Ltd. (Seoul, Korea). The challenge strain, *M. tuberculosis* H37Ra, was purchased from the American Type Culture Collection (ATCC 25177, VA, USA). These strains were cultured at 37°C with agitation at 100–200 rpm in Middlebrook 7H9 broth (Difco, MI, USA) supplemented with 0.2% (v/v) glycerol (Sigma-Aldrich, Gillingham, UK), 10% albumin-dextrose-catalase (ADC), and 0.05% (v/v) Tween 80 (Sigma-Aldrich, Gillingham, UK). The incubation period was more than 2 weeks for the H37Ra strain and 1–2 days for rMs. Following incubation, the bacterial cultures were subjected to glass bead vortexing and passed through a 25-gauge needle. The supernatant was harvested after centrifugation at 3,300 × g for 10 min, and the collected bacterial pellets were resuspended in 10% glycerol and stored at or below -80°C until use. Middlebrook 7H10 agar (Millipore, Burlington, MA, USA) was used as the solid medium, prepared by mixing 10 g of agar base, 2.5 mL of glycerol, and 10 mL of oleic acid-albumin-dextrose-catalase (OADC; Becton Dickinson, Sparks, MD, USA) per 500 mL of distilled water. After sterilization, 25–30 mL of the medium was poured into each Petri dish (SPL Life Sciences, Pocheon, Korea) and stored at 4°C prior to use.

**Fig 1.**
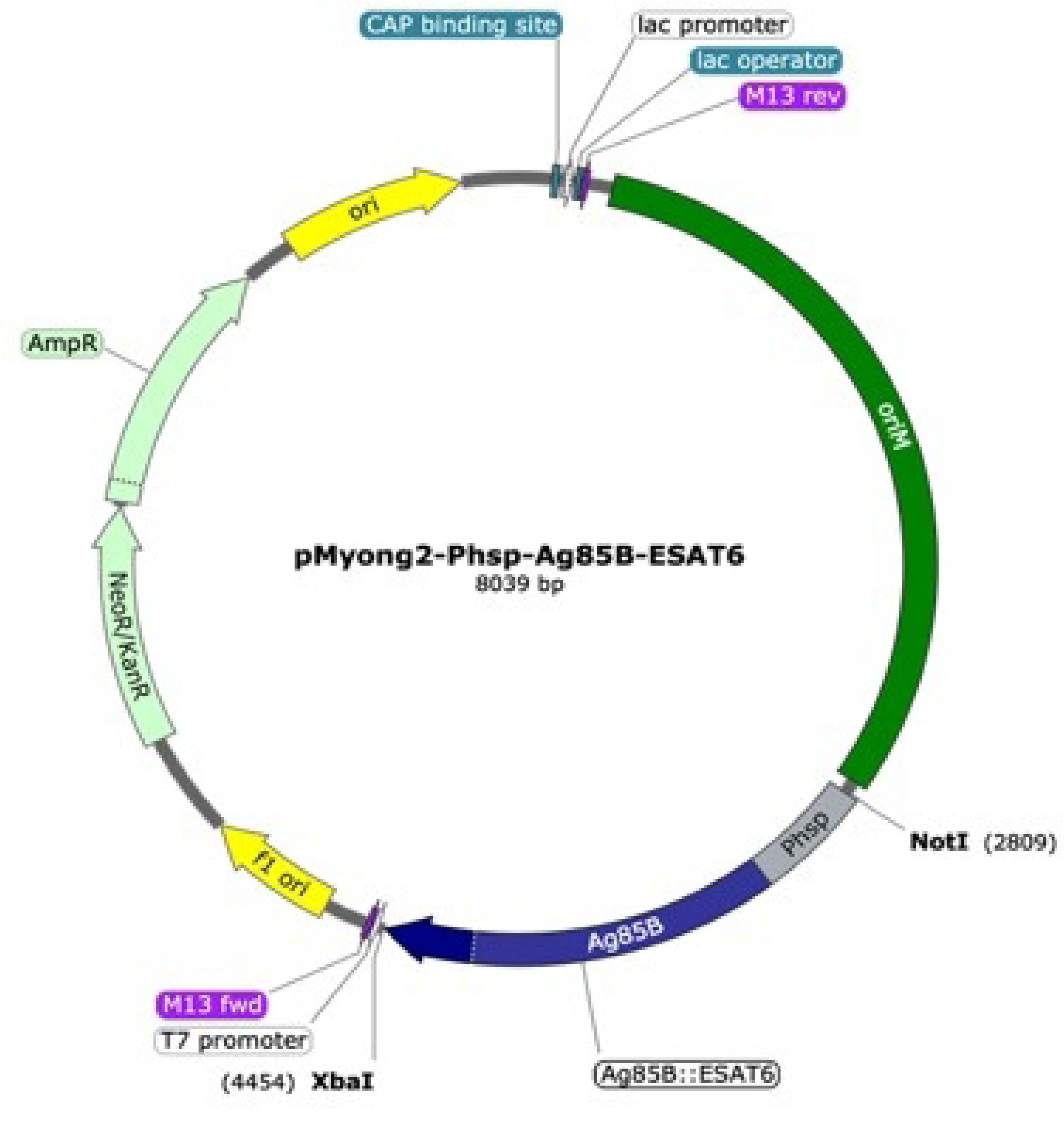
Construction of the recombinant pMyong2-Phsp-Ag85B-ESAT-6 expression vector. Schematic map of the pMyong2-based recombinant plasmid encoding the *Mycobacterium tuberculosis (M. tuberculosis)* antigens Ag85B and ESAT-6 under the control of the mycobacterial heat shock protein (*hsp*) promoter. The construct was designed for expression in *Mycobacterium smegmatis* (*M. smegmatis*) as a live recombinant vaccine candidate to enhance BCG-induced protective immunity against *M. tuberculosis*.

### Vaccines, Immunization & Challenge

Animal experiments were conducted using 4-week-old female BALB/c mice infected with the *M.. tuberculosis* H37Ra strain under Biosafety Level 2 (BSL-2) conditions, based on previously described protocols (11–13). Except for the negative control group, all groups received a single subcutaneous injection of 0.1 mL of a commercially available BCG vaccine (*Mycobacterium bovis* Danish 1331 strain, 2.0 x 10^6^ - 8.0 x 10^6^ colony-forming units (CFUs) /mL). Booster vaccinations were administered twice in total. The group receiving two additional doses of the same commercial BCG vaccine was designated as BCG 2 group, while other booster groups were administered two doses of the rMs vaccine at varying doses, either alone or combined with β-glucan (6 mg/mL) extracted and purified from fermented oats from CLIPS BnC Co., Ltd. (Seoul, Korea) (Fig. 2b). The detailed immunization schedule and experimental groups are presented in Figures 2a and 2b.

**Fig 2.**
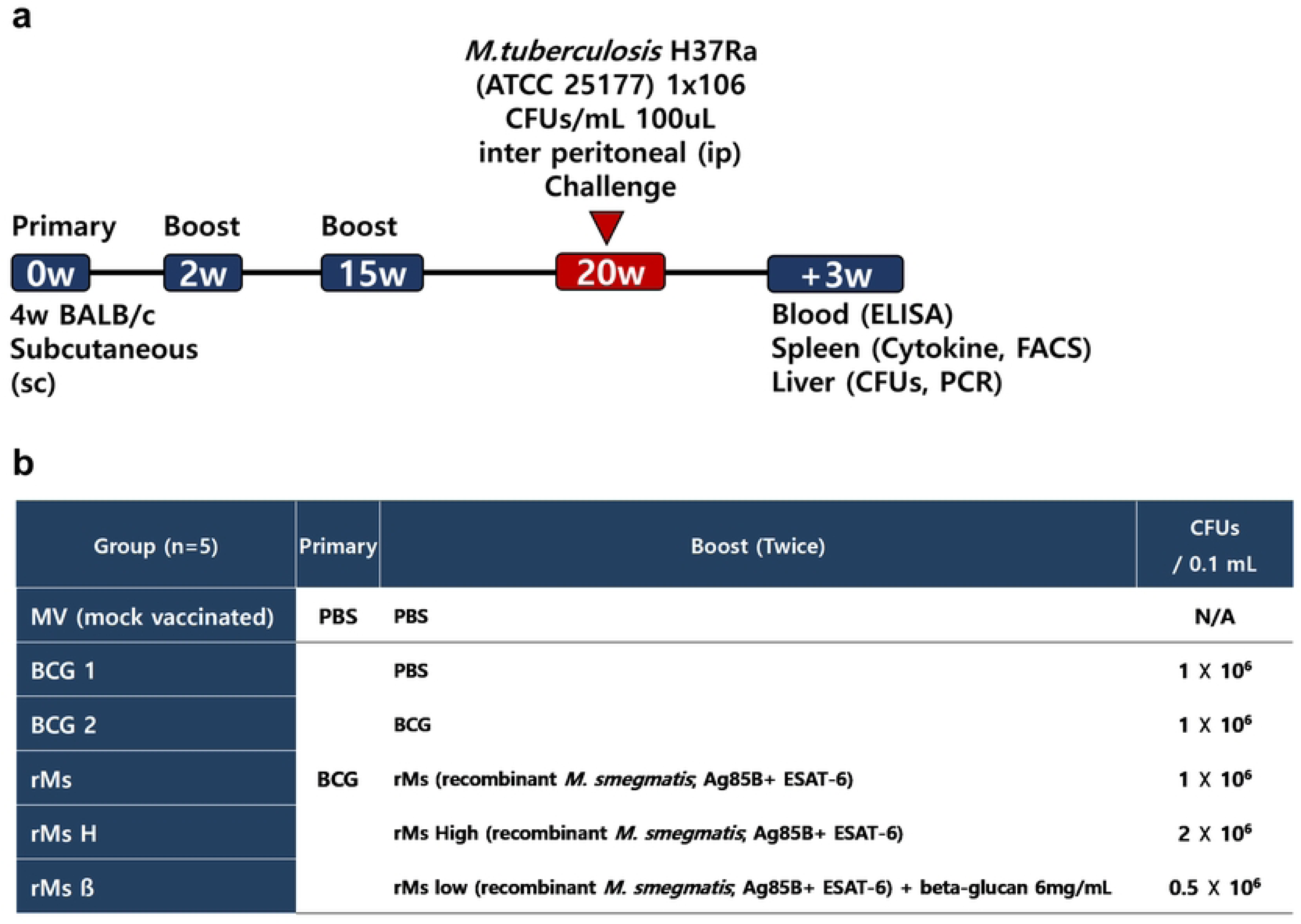
Study schedule & study group. **(a)** Four-week-old female BALB/c mice were vaccinated subcutaneously a total of three times. Five weeks following the final immunization, mice were challenged via intraperitoneal injection with 100 μL of *M. tuberculosis* H37Ra (ATCC 25177) suspension in PBS 1×10^6^ CFUs /mL (OD_600_ = 0.1). Three weeks post-challenge, blood, spleen and liver tissues were harvested for ELISA, cytokine analysis, flow cytometry, and colony counting assays. **(b)** Mice were divided into six experimental groups (n = 5). Control groups included a mock-vaccinated negative control group receiving PBS only (MV), a single-BCG primed group boosted with PBS (BCG 1), and a group receiving a primary BCG dose followed by two BCG booster vaccinations (BCG 2). The experimental booster groups received a recombinant *M. smegmatis* vector expressing Ag85B and ESAT-6 antigens at a standard dose of 1×10^6^ CFUs (rMs), a high dose of 2×10^6^ CFUs (rMs H), or a low dose of 0.5×10^6^ CFUs formulated with 6 mg/mL β-glucan extract (rMs β).

Five weeks after the final vaccination, the mice were challenged via intraperitoneal injection with 100μL of *M. tuberculosis* H37Ra (ATCC 25177). Three weeks post-challenge, blood, liver and spleen tissues were harvested to evaluate immunogenicity (Fig. 2b). All animal procedures were approved by the Institutional Animal Care and Use Committee (IACUC) of the School of Medicine, The Catholic University of Korea (Approval No. CUMC-2021-0036-01) and performed at the Laboratory Animal Research Center of The Catholic University of Korea in strict compliance with the Animal Welfare Act and the Laboratory Animal Act. All experiments, including animal studies, were conducted at the Vaccine Bio Research Institute, College of Medicine, The Catholic University of Korea.

### Bacterial Clearance Analysis

Mouse challenge experiments were performed based on previously reported protocols (14–16). Following intraperitoneal administration of the *M. tuberculosis* H37Ra strain to BALB/c mice, survival rates and abnormal clinical signs were monitored for 8 days. Three weeks post-challenge, liver tissues were harvested from the mice and homogenized in 1 mL of PBS containing 1% Tween-80. A 400μL aliquot of the homogenate was spread onto Middlebrook 7H10 agar, incubated at 37°C for 10 days, and the CFUs were enumerated. In parallel, naive BALB/c mice were challenged using the same procedure, and their liver tissues were harvested to extract total cellular DNA using the QIAamp DNA Mini Kit (Qiagen, Hilden, Germany) according to the manufacturer’s instructions. Using the extracted DNA as a template, PCR was performed with IS6110-specific primers (Table 1) (17, 18) to identify the *in vivo* persistence of the H37Ra strain.

**Table 1.**
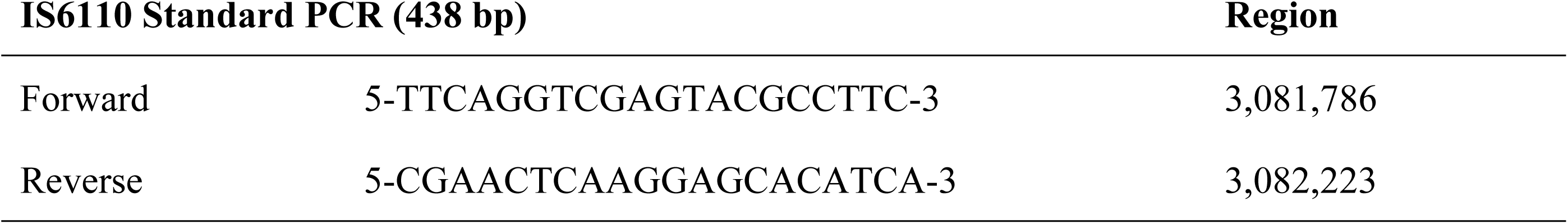
IS6100 primers for H37Ra strain detecting PCR.

### Flow Cytometry Analysis

For flow cytometric analysis, single-cell suspensions were prepared from the spleen, and 20,000 cells per mouse were harvested. Cells were stimulated for 3 days with ESAT-6 (0.5μg/mL) or Ag85B (1μg/mL) in the presence of GolgiPlug (BD Biosciences, San Jose, CA, USA). Live/Dead staining was performed using Fixable Viability Stain 700 (BD Biosciences, San Jose, CA, USA). To minimize non-specific antibody binding, Fc receptors were blocked using anti-mouse CD16/CD32 antibody (BD Biosciences, San Jose, CA, USA). Subsequently, cell surface staining was conducted using anti-mouse CD4-FITC, CD3-PE-Cy7, and CD8-PerCP-Cy5.5 antibodies (BD Biosciences, San Jose, CA, USA). After cell fixation using Cytofix (BD Biosciences, San Jose, CA, USA), intracellular cytokine staining was performed using anti-mouse IFN-γ -APC and IL-17A-BV421 antibodies. Fluorescence was measured using a FACS Aria Fusion flow cytometer (BD Biosciences, San Jose, CA, USA), and data were analyzed using FlowJo software version 10.10.0 (BD Life Sciences, Ashland, OR, USA).

### Cytokine ELISA

Three weeks post-immunization, single-cell suspensions were prepared from the spleens of five mice per group and stimulated with ESAT-6 (0.5 μg/mL; Sino Biological, Beijing, China) or Ag85B (1 μg/mL; Sino Biological) in the presence of GolgiPlug (BD Biosciences, San Jose, CA, USA) for three days. Following incubation, the cell culture supernatants were harvested, and the concentrations of IFN-γ and IL-17A were quantified using commercial cytokine ELISA kits (Proteintech, Rosemont, IL, USA) according to the manufacturer’s instructions.

### Humoral Immune Responses to ESAT-6 and Ag85B

To determine ESAT-6- and Ag85B-specific antibody titers, an enzyme-linked immunosorbent assay (ELISA) was performed. Briefly, 96-well ELISA plates (SPL Life Sciences, Pocheon, Korea) were coated with ESAT-6 (0.5 μg/mL; Sino Biological, Beijing, China) or Ag85B (1 μg/mL; Sino Biological) dissolved in 0.05 M carbonate/bicarbonate coating buffer (pH 9.6) and incubated overnight at 4 °C. After washing with phosphate-buffered saline (PBS) containing 0.05% (v/v) Tween 20 (PBST; Sigma-Aldrich, Gillingham, UK), the plates were blocked with 5% skim milk in PBS for 1 h at room temperature. Serum samples were serially diluted 10-fold starting from an initial dilution of 1:8, added to each well, and incubated for 1.5 h at room temperature. Following wash steps, Horseradish Peroxidase(HRP)-conjugated secondary antibodies—goat anti-mouse IgG (1:100,000; Invitrogen/Novex), goat anti-mouse IgG2c (1:10,000; Invitrogen, Carlsbad, CA, USA), and goat anti-mouse IgA (1:4,000; Invitrogen)—were added and incubated. After additional washing, the color reaction was developed by adding 3,3’,5,5’-tetramethylbenzidine (TMB) substrate and terminated with 0.8 M H₂SO₄. Absorbance was measured at 450 nm using a SpectraMax iD3 microplate reader (Molecular Devices, San Jose, CA, USA). ESAT-6- and Ag85B-specific antibody levels were evaluated and compared between groups (n = 5 per group) using optical density (OD₄₅₀) values obtained at a 1:8 serum dilution.

### Statistical analyses

Statistical differences among groups were evaluated by one-way analysis of variance (ANOVA) followed by Tukey’s multiple comparison test using GraphPad Prism software (version 10.0; GraphPad Software, San Diego, CA, USA). Different significance levels were designated as * *p* < 0.05, ** *p* < 0.01, *** *p* < 0.001, and **** *p* < 0.0001.

## Results

### Survival rates & Bacterial clearance

Mice were monitored daily for survival and abnormal clinical signs—including inflammation spread at the site of inoculation or challenge, ulceration, sudden body weight loss exceeding 20%, severe anorexia lasting more than 3 days, limb paralysis, severe ataxia, and bone fractures— for 8 days post-challenge; no notable abnormalities were observed across all groups. However, on day 1 post-challenge, one mouse died in each of the rMs and rMs H groups, resulting in an 80% survival rate while all other groups maintained a 100% survival rate (Fig. 3a).

**Figure 3.**
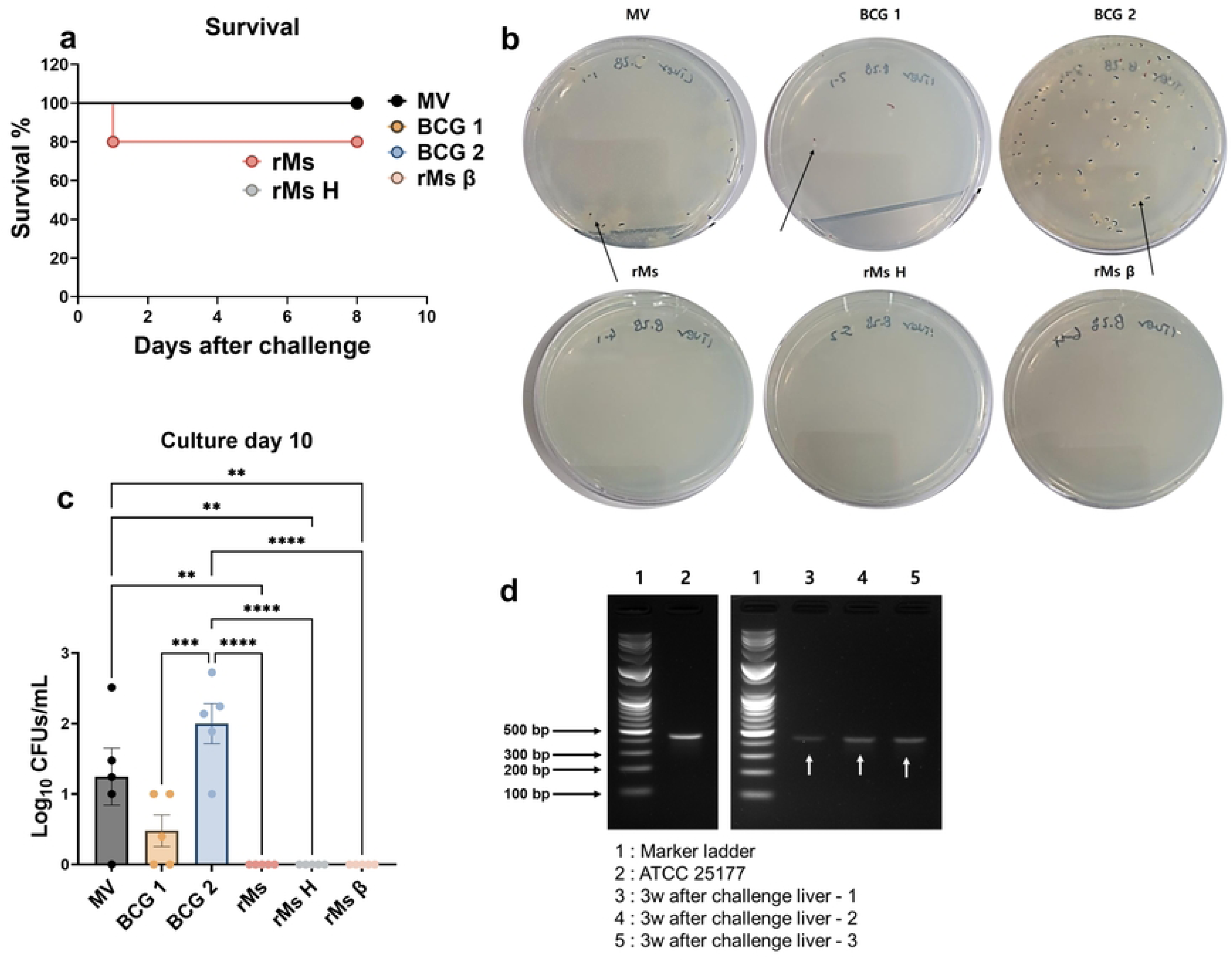
Results of survival and bacterial clearance. **(a)** Mice were monitored daily for survival and clinical signs for 8 days post-challenge. **(b, c)** Three weeks post-challenge, liver tissues (left major and minor lobes) were harvested and homogenized in 1 mL of PBS containing 1% Tween 20. A 400 μL aliquot of the homogenate was plated onto Middlebrook 7H10 agar and incubated for 10 days to quantify bacterial colonies (n = 5 per group; n = 4 for rMs and rMs H groups). Bacterial counts were log₁₀-transformed and presented as mean ± standard error of the mean (SEM). Statistical significance among groups was determined by one-way ANOVA with Tukey’s multiple comparison test (* *p* < 0.05, ** *p* < 0.01, *** *p* < 0.001, and **** *p* < 0.0001). **(d)** Normal mice (n = 3) were intraperitoneally challenged with the *Mycobacterium tuberculosis* H37Ra strain at a dose of 1 × 10⁵ CFU. Three weeks post-challenge, genomic DNA was extracted from liver tissues, and PCR was performed using IS6100-specific primers to verify an expected band at 438 bp.

No bacterial colonies were detected up to day 10 of culture in both the rMs-immunized group and the low-dose combined with β-glucan group, showing a statistically significant decrease compared to the MV (** *p* < 0.01) and BCG 2 groups (**** *p* < 0.0001) (Fig. 3b, 3c). The BCG 2 group, which received a booster dose of BCG vaccine, exhibited a significantly higher colony count at day 10 of culture compared to all other groups except the MV group (Fig. 3b, 3c). Notably, the BCG 2 group showed a significantly higher bacterial load than the single-dose BCG 1 group (*p* = 0.0008). The log_₁₀_ CFU count was 1.246 for the MV group and 1.999 for the BCG 2 group, representing a 1.6-fold higher CFU value in the BCG 2 group compared to the MV group (Fig. 3c). Complete survival (100%) and the absence of bacterial colonies were observed exclusively in the low-dose combined with β-glucan group (rMs β). To confirm normal infection with the *Mycobacterium tuberculosis* strain H37Ra, PCR was performed using IS6100 primers, a representative marker for tuberculosis identification (17, 18). PCR amplification using DNA extracted from the H37Ra strain and the liver tissue of a normal mouse infected with *M. tuberculosis* strain H37Ra both yielded a clear band at 438 bp, confirming that intraperitoneal administration of H37Ra successfully induced liver infection (Fig. 3d).

### T cell & Cytokine responses

Regarding T cells responses derived from splenocyte single-cell suspensions, stimulation with the Ag85B antigen induced prominent expression of both IFN-γ (*p* =0.0043) and IL-17A (*p* =0.0093) in the rMs group, showing statistically significant differences compared to the MV group. Furthermore, for IFN-γ expression, a statistically significant difference was also observed between the rMs group and the rMs β group (*p* = 0.0032). Across all groups, CD4⁺ T cells accounted for more than 50% of CD3⁺ T cells, with no significant differences observed among the groups. The expression levels of IFN-γ⁺ and IL-17A⁺ in T cells were calculated as percentages relative to CD4⁺ T cells; no significant differences among groups were observed under conditions other than Ag85B antigen stimulation (Fig. 4a, 4c). In the cytokine analysis of total splenocyte single-cell suspensions, the rMs, rMs β, and BCG 1 groups exhibited high responsiveness, although no statistically significant differences were observed among the groups. For IFN-γ release, the highest levels were detected in the rMs group under media control conditions (6,383 pg/mL), the rMs β group upon ESAT-6 stimulation (6,571 pg/mL), and the BCG 2 group upon Ag85B stimulation (4,292 pg/mL). For IL-17A, the highest cytokine responses were recorded in the rMs β group under media control conditions (4,227 pg/mL), and in the BCG 2 group upon both ESAT-6 (4,170 pg/mL) and Ag85B (2,111 pg/mL) stimulations (Fig. 4b, 4d).

**Figure 4.**
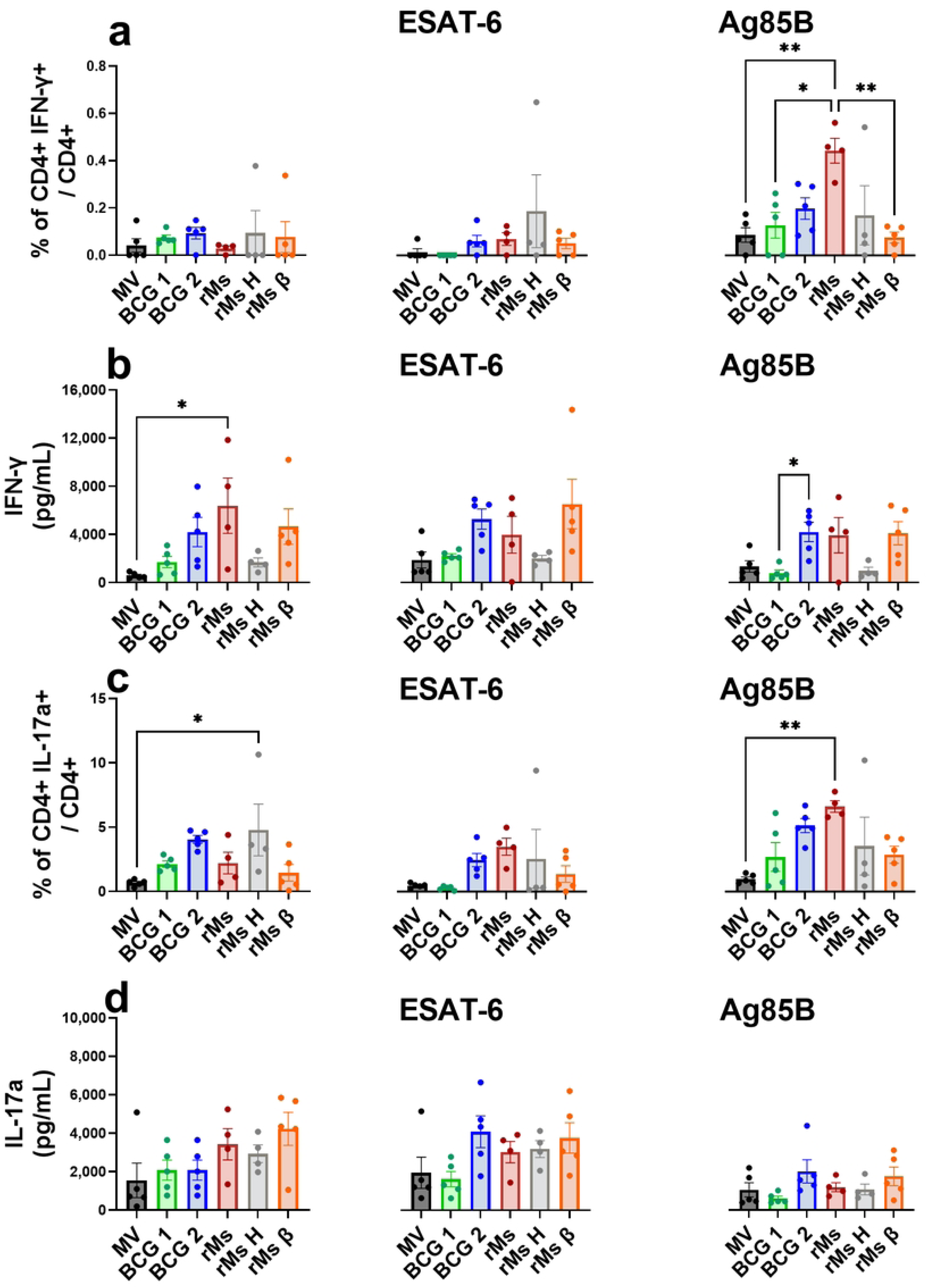
Results of Cytokine & T cell responses. Cytokine concentrations in culture supernatants of mouse splenocyte single-cell suspensions were quantified using commercial ELISA kits. To evaluate cytokine expression in CD4⁺ T cells, 20,000 cells per sample were harvested and stained with primary antibodies against CD3, CD4, CD8, IFN-γ, and IL-17a (n = 5 per group; n = 4 for rMs and rMs H groups). Cytokine levels are expressed in pg/mL, and cytokine-producing T cells are presented as percentages relative to total CD4⁺ T cells. IFN-γ expression was evaluated by **(a)** the proportion of CD4⁺IFN-γ⁺ cells among CD4⁺ T cells (%) and **(b)** ELISA quantification of secreted IFN-γ in culture supernatants. IL-17a^+^ expression was determined by **(c)** the proportion of CD4⁺IL-17a^+^ cells among CD4⁺ T cells (%) and **(d)** ELISA quantification of secreted IL-17a^+^ in culture supernatants. All data are presented as mean ± SEM. Statistical differences among groups were analyzed by one-way ANOVA with Tukey’s multiple comparison test (* p < 0.05, ** p < 0.01, *** p < 0.001, and **** p < 0.0001).

### Antibody response for ESAT-6 & Ag85B antigen

Antibody responses were evaluated based on absorbance at OD_450_ using serum samples collected three weeks post-challenge (Fig. 5). The ESAT-6-specific antibody responses were determined for IgG, IgG2c, and IgA subtypes. For IgG, the rMs H group exhibited the highest antibody response, with a mean OD₄₅₀ value of 1.35, showing statistically significant differences compared to both the BCG 1 (*p* =0.0015) and rMs groups (*p* = 0.0034). For the IgA subtype, the BCG 2 group demonstrated the highest mean OD₄₅₀ value of 0.44, which was statistically significantly different from all other experimental groups (** *p* < 0.01) except the MV group (Fig. 5a, 5b, 5c). Regarding Ag85B-specific antibody responses, the BCG 2 group showed the highest levels across all Ig subtypes; however, no statistically significant differences were observed among the groups. In the BCG 2 group, the mean OD₄₅₀ values were 1.63 for IgG, 0.41 for IgG2c, and 0.53 for IgA (Fig. 5d, 5e, 5f).

**Figure 5.**
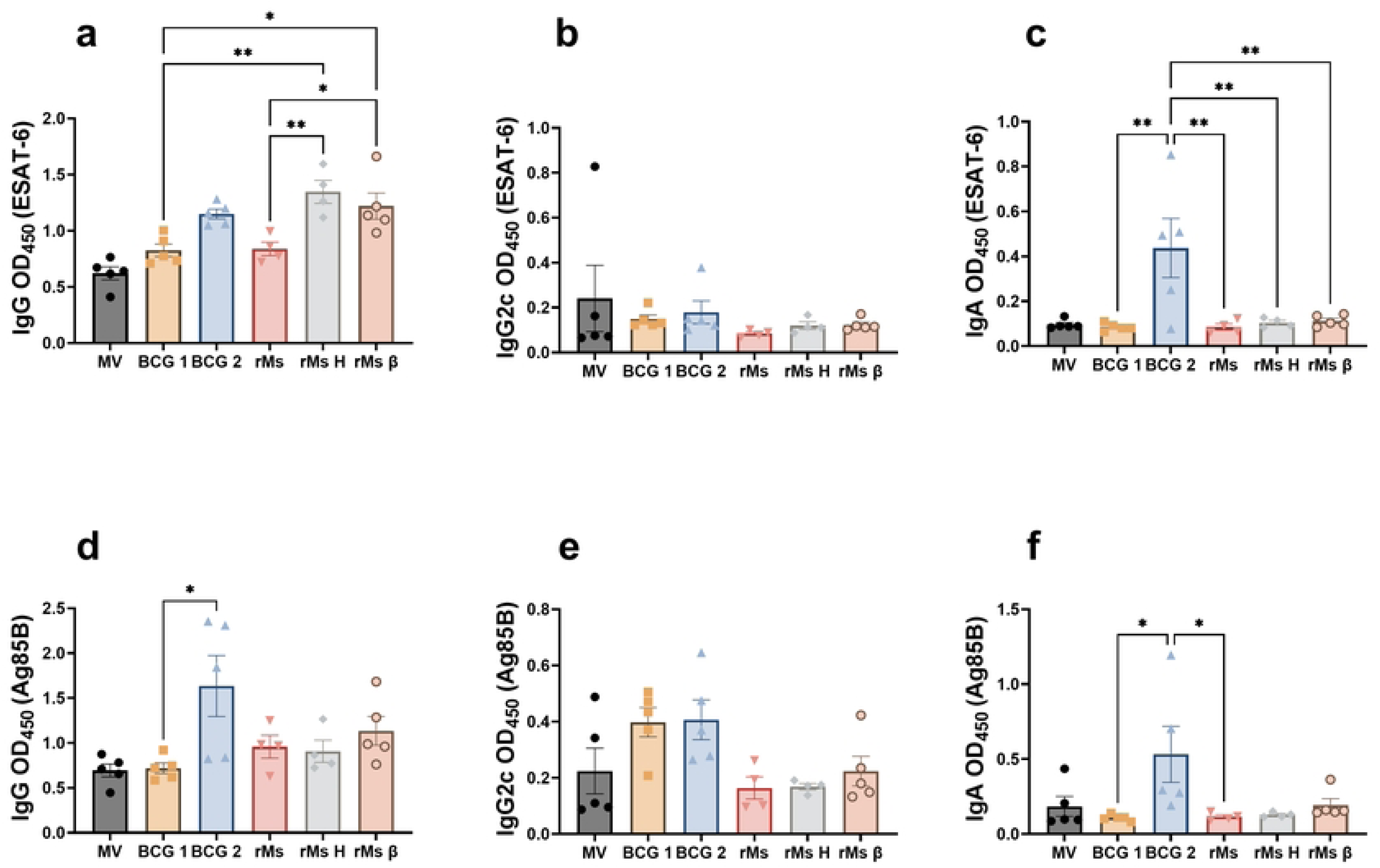
Antibody responses for ESAT-6 and Ag85B. ELISA plates were coated with recombinant **(a, b, c)** ESAT-6 (0.5 μg/mL) or **(d, e, f)** Ag85B (1 μg/mL) proteins (Sino Biological, Beijing, China). Antigen-specific IgG, IgG2c, and IgA antibody responses were evaluated by measuring absorbance at 450 nm (OD₄₅₀). All OD₄₅₀ values are presented as mean ± standard error of the mean (SEM). Statistical differences among groups were determined by one-way ANOVA with Tukey’s multiple comparison test (*p < 0.05, ** p < 0.01, *** p < 0.001, and **** p < 0.0001).

## Discussion

Although BCG effectively prevents severe forms of tuberculosis in infants and young children, its protective efficacy against pulmonary tuberculosis in adolescents and adults remains inconsistent (4, 19, 20). To overcome these limitations, various next-generation tuberculosis vaccine candidates— including recombinant BCG, attenuated *M. tuberculosis* strains, protein subunit vaccines, and viral vector-based vaccines—are currently under development. Among these, M72/AS01E, an adjuvanted protein subunit vaccine, is considered one of the most promising candidates to date, demonstrating approximately 50% protective efficacy in clinical trials involving adults with LTBI (21, 22). However, high-level protection was not achieved across all target populations, highlighting the need for further validation regarding long-term durability and broader applicability (21, 22). Furthermore, although live attenuated candidates such as recombinant BCG (VPM1002) and attenuated *M. tuberculosis* (MTBVAC) are undergoing clinical evaluation, superior clinical efficacy that reliably replaces BCG for adult pulmonary tuberculosis has yet to be definitively established (23, 24).

One of the major hurdles in tuberculosis vaccine development is that robust immunogenicity does not necessarily translate into protective efficacy. For example, the viral vector-based vaccine candidate MVA85A induced strong Ag85B-specific CD4⁺ T-cell responses but failed to demonstrate significant protective efficacy in a large-scale clinical trial (11, 25), highlighting the continued difficulty in defining reliable immune correlates of protection against tuberculosis (12, 13). Against this background, we developed an rMs booster co-expressing Ag85B and ESAT-6 using the pMyong2 expression vector and evaluated oat-derived β-glucan as an adjuvant to enhance BCG-primed protection.

In this study, we employed an *M. tuberculosis* H37Ra challenge model, an attenuated strain that can be safely handled under Biosafety Level 2 (BSL-2) conditions (26). Previous studies have demonstrated that H37Ra can establish infection and disseminate to extra pulmonary organs, including the liver and spleen, in BALB/c mice (27, 28), and our preliminary experiments similarly confirmed stable bacterial detection in the liver following intraperitoneal challenge (Fig. 3b). However, representing the most critical limitation of this study, the stringency and translational relevance of this model should be considered in the context of contemporary preclinical TB vaccine studies, which commonly employ virulent *M. tuberculosis*, particularly low-dose aerosol H37Rv challenge, and assess bacterial burdens in the lungs and spleen together with pulmonary pathology. For example, BCG-primed mice boosted with the whole-cell DAR-901 vaccine were evaluated following low-dose aerosol H37Rv challenge using lung and spleen CFUs as efficacy endpoints (29), while recombinant *M. smegmatis* expressing Ag85B-ESAT6 has also been evaluated against virulent H37Rv using lung bacterial burden and pulmonary pathology (30). More recent studies have further demonstrated that ultra-low-dose aerosol infection initiated with only a few bacilli may more closely reproduce important features of heterogeneous human pulmonary TB (31). Therefore, the intraperitoneal H37Ra/liver-CFU model used in the present study should be regarded as a lower-stringency surrogate model for the preliminary evaluation of systemic mycobacterial clearance rather than a direct model of protection against pulmonary infection with virulent *M. tuberculosis*. Accordingly, efficacy outcomes obtained using this model should be interpreted as evidence of systemic mycobacterial control within this experimental setting rather than as evidence of sterilizing immunity or equivalent protection against virulent H37Rv aerosol challenge.

In preclinical tuberculosis vaccine studies, vaccine efficacy is commonly evaluated by quantifying bacterial loads as CFUs in the lungs, spleen, or liver following M. tuberculosis challenge. While most studies report vaccine efficacy in terms of log reduction in tissue bacterial burdens, the absence of detectable CFUs in target organs remains relatively uncommon (20, 32–34). In this regard, no detectable liver CFUs were observed following H37Ra challenge in groups receiving BCG priming followed by rMs, rMs H, or rMs β boosting (Fig. 3c, 3d). Notably, the rMs β group achieved a 100% survival rate (Fig. 3a) despite receiving only half the dose (5 × 10⁵ CFUs) compared to the standard rMs group (1 × 10⁶ CFUs) (Fig. 2b), underscoring the value of further investigation to elucidate the precise immunological mechanisms involved. Conversely, revaccination with the commercially available BCG vaccine yielded lower bacterial clearance compared to a single BCG dose, consistent with previous reports demonstrating that repeated BCG vaccination fails to provide a clear booster effect (Fig. 3c, 3d) (19, 35, 36). Collectively, these findings indicate that the rMs-based heterologous booster strategy enhanced mycobacterial clearance in the H37Ra challenge model compared with homologous BCG revaccination.

These findings are consistent with previous studies supporting recombinant *M. smegmatis* as an immunogenic vaccine vector and with studies evaluating Ag85B- and ESAT-6-based tuberculosis vaccines. Previous studies have reported that recombinant *M. smegmatis* enhances Th1 immune responses and can induce protective activity against M. tuberculosis (37, 38). Furthermore, Ag85B- and ESAT-6-based vaccines have been shown to boost antigen-specific CD4⁺ T-cell responses and reduce bacterial burden following BCG priming (39–42). Consistently, the rMs group in this study exhibited robust Ag85B- and ESAT-6-specific IFN-γ- and IL-17A-expressing CD4⁺ T-cell responses (Fig. 4), which may have contributed to the enhanced mycobacterial clearance observed after H37Ra challenge. Additionally, given reports that recombinant *M.Smegmatis* can promote macrophage activation, phagosome maturation, and intracellular bactericidal mechanisms (43), the potential involvement of innate immune activation driven by the vector strain itself cannot be excluded.

However, as the second major limitation of this study, our findings also showed that the measured immunological responses did not consistently predict bacterial clearance. Robust antigen-specific Th1/Th17 responses were most evident in the rMs group, whereas the rMs β and rMs H groups showed more limited antigen-specific responses despite similarly undetectable liver CFUs following H37Ra challenge (Fig. 4). Conversely, the BCG 2 group, which showed the poorest bacterial control, displayed CD4⁺ T-cell and cytokine responses comparable to those of the rMs β and rMs H groups, together with elevated IgA responses (Fig. 5). These findings indicate that the antigen-specific T-cell and antibody responses measured here are insufficient to explain the differences in bacterial control among groups. This pattern is consistent with the broader difficulty in defining reliable correlates of protection for tuberculosis vaccines, as illustrated by MVA85A, in which strong vaccine-induced cellular immune responses did not translate into significant protective efficacy (22–25, 44).

## Conclusion

In summary, a heterologous BCG prime–boost strategy using pMyong2-based recombinant *M. smegmatis* (rMs) expressing Ag85B and ESAT-6 superiorly enhanced mycobacterial clearance in an H37Ra challenge model compared to BCG revaccination. Furthermore, oat-derived β-glucan served as an effective dose-sparing adjuvant, allowing a 50% dose reduction without compromising protective efficacy. Despite incomplete correlation with measured adaptive immune parameters, these results position rMs as a viable heterologous booster candidate, warranting further mechanistic investigation and validation in virulent pulmonary challenge models.

## List of abbreviations

BCG: Bacille Calmette–Guérin
BCG 1: a single-BCG primed group boosted with PBS
BCG 2: a group receiving a primary BCG dose followed by two BCG booster vaccinations
CFUs: Colony forming units
*hsp*: heat shock protein
LTBI: latent tuberculosis infection
MDR-TB: multidrug-resistant TB
MV: mock-vaccinated negative control group
M. smegmatis: Mycobacterium smegmatis
M. tuberculosis: Mycobacterium tuberculosis
NTM: non-tuberculous mycobacteria
rMs: booster groups received a recombinant *M. smegmatis* vector expressing Ag85B and ESAT-6 antigens at a standard dose of 1×10^6^ CFUs (rMs)
rMs H: booster groups received a recombinant *M. smegmatis* vector expressing Ag85B and ESAT-6 antigens at a high dose of 2×10^6^ CFUs (rMs H)
rMs β: booster groups received a recombinant *M. smegmatis* vector expressing Ag85B and ESAT-6 antigens at a low dose of 0.5×10^6^ CFUs formulated with 6 mg/mL β-glucan extract
SEM: standard error of the mean
TB: Tuberculosis

## Acknowledgments

Not applicable.

## Data Availability Statement

The datasets used and/or analyzed during the current study are available from the corresponding author upon reasonable request.

## References

1. World Health Organization. Global tuberculosis report 2024 [Internet]. Geneva: World Health Organization; 2024 [cited 2026 Jul 22] 68 p. (Available from: https://www.who.int/teams/global-programme-on-tuberculosis-and-lung-health/tb-reports/global-tuberculosis-report-2024)

2. Furin J, Cox H, Pai M. Tuberculosis. The Lancet. 2019;393(10181):1642–56. [10.1016/S0140-6736(19)30308-3]

3. World Health Organization. BCG vaccines: WHO position paper – February 2018. Wkly Epidemiol Rec. 2018 Feb 23;93 (8) :73–96. (Available from: https://www.who.int/publications/i/item/who-wer9308-73-96

4. Mangtani P, Abubakar I, Ariti C, Beynon R, Pimpin L, Fine PE, et al. Protection by BCG vaccine against tuberculosis: a systematic review of randomized controlled trials. Clin Infect Dis. 2014;58(4):470–80. [10.1093/cid/cit790]

5. Colditz GA, Brewer TF, Berkey CS, Wilson ME, Burdick E, Fineberg HV, et al. Efficacy of BCG vaccine in the prevention of tuberculosis. Meta-analysis of the published literature. Jama. 1994;271(9):698–702. [10.1001/jama.1994.03510330076038]

6. Fine PE. Variation in protection by BCG: implications of and for heterologous immunity. Lancet. 1995;346(8986):1339–45. [10.1016/s0140-6736(95)92348-9]

7. Shah JA, Lindestam Arlehamn CS, Horne DJ, Sette A, Hawn TR. Nontuberculous Mycobacteria and Heterologous Immunity to Tuberculosis. J Infect Dis. 2019;220(7):1091–8. [10.1093/infdis/jiz285]

8. Schrager LK, Harris RC, Vekemans J. Research and development of new tuberculosis vaccines: a review. F1000Res. 2018;7:1732. [10.12688/f1000research.16521.2]

9. Brennan MJ, Thole J. Tuberculosis vaccines: a strategic blueprint for the next decade. Tuberculosis (Edinb). 2012;92 Suppl 1:S6–13. [10.1016/s1472-9792(12)70005-7]

10. Lee H, Kim BJ, Kim BR, Kook YH, Kim BJ. The development of a novel Mycobacterium-Escherichia coli shuttle vector system using pMyong2, a linear plasmid from Mycobacterium yongonense DSM 45126T. PLoS One. 2015;10(3):e0122897. [10.1371/journal.pone.0122897]

11. Tameris M, Geldenhuys H, Luabeya AK, Smit E, Hughes JE, Vermaak S, et al. The candidate TB vaccine, MVA85A, induces highly durable Th1 responses. PLoS One. 2014;9(2):e87340. [10.1371/journal.pone.0087340]

12. McShane H, Williams A. A review of preclinical animal models utilised for TB vaccine evaluation in the context of recent human efficacy data. Tuberculosis (Edinb). 2014;94(2):105–10. [10.1016/j.tube.2013.11.003]

13. Kaufmann SHE, Dockrell HM, Drager N, Ho MM, McShane H, Neyrolles O, et al. TBVAC2020: Advancing Tuberculosis Vaccines from Discovery to Clinical Development. Front Immunol. 2017;8:1203. [10.3389/fimmu.2017.01203]

14. Garcia-Pelayo MC, Bachy VS, Kaveh DA, Hogarth PJ. BALB/c mice display more enhanced BCG vaccine induced Th1 and Th17 response than C57BL/6 mice but have equivalent protection. Tuberculosis (Edinb). 2015;95(1):48–53. [10.1016/j.tube.2014.10.012]

15. Hanna CC, Ashhurst AS, Quan D, Maxwell JWC, Britton WJ, Payne RJ. Synthetic protein conjugate vaccines provide protection against Mycobacterium tuberculosis in mice. Proc Natl Acad Sci U S A. 2021;118(4). [10.1073/pnas.2013730118]

16. Kadir NA, Sarmiento ME, Acosta A, Norazmi MN. Cellular and humoral immunogenicity of recombinant Mycobacterium smegmatis expressing Ag85B epitopes in mice. Int J Mycobacteriol. 2016;5(1):7–13. [10.1016/j.ijmyco.2015.09.006]

17. El Khéchine A, Henry M, Raoult D, Drancourt M. Detection of Mycobacterium tuberculosis complex organisms in the stools of patients with pulmonary tuberculosis. Microbiology (Reading). 2009;155(Pt 7):2384–9. [10.1099/mic.0.026484-0]

18. Thierry D, Brisson-Noël A, Vincent-Lévy-Frébault V, Nguyen S, Guesdon JL, Gicquel B. Characterization of a Mycobacterium tuberculosis insertion sequence, IS6110, and its application in diagnosis. J Clin Microbiol. 1990;28(12):2668–73. [10.1128/jcm.28.12.2668-2673.1990]

19. Kaufmann SHE. Vaccination Against Tuberculosis: Revamping BCG by Molecular Genetics Guided by Immunology. Front Immunol. 2020;11:316. 10.3389/fimmu.2020.00316

20. Andersen P, Scriba TJ. Moving tuberculosis vaccines from theory to practice. Nat Rev Immunol. 2019;19(9):550–62. [10.1038/s41577-019-0174-z]

21. Van Der Meeren O, Hatherill M, Nduba V, Wilkinson Robert J, Muyoyeta M, Van Brakel E, et al. Phase 2b Controlled Trial of M72/AS01E Vaccine to Prevent Tuberculosis. New England Journal of Medicine. 2018;379(17):1621–34. [10.1056/nejmoa1803484]

22. Tait Dereck R, Hatherill M, Van Der Meeren O, Ginsberg Ann M, Van Brakel E, Salaun B, et al. Final Analysis of a Trial of M72/AS01E Vaccine to Prevent Tuberculosis. New England Journal of Medicine. 2019;381(25):2429–39. [10.1056/nejmoa1909953]

23. Nieuwenhuizen NE, Kulkarni PS, Shaligram U, Cotton MF, Rentsch CA, Eisele B, et al. The Recombinant Bacille Calmette-Guérin Vaccine VPM1002: Ready for Clinical Efficacy Testing. Front Immunol. 2017;8:1147. [10.3389/fimmu.2017.01147]

24. Lacámara S, Martin C. MTBVAC: A Tuberculosis Vaccine Candidate Advancing Towards Clinical Efficacy Trials in TB Prevention. Arch Bronconeumol. 2023. [10.1016/j.arbres.2023.09.009]

25. Tameris MD, Hatherill M, Landry BS, Scriba TJ, Snowden MA, Lockhart S, et al. Safety and efficacy of MVA85A, a new tuberculosis vaccine, in infants previously vaccinated with BCG: a randomised, placebo-controlled phase 2b trial. Lancet. 2013;381(9871):1021–8. [10.1016/s0140-6736(13)60177-4]

26. Lenaerts A, Barry CE, 3rd, Dartois V. Heterogeneity in tuberculosis pathology, microenvironments and therapeutic responses. Immunol Rev. 2015;264(1):288–307. [10.1111/imr.12252]

27. Williams A, Orme IM. Animal Models of Tuberculosis: An Overview. Microbiol Spectr. 2016;4(4). [10.1128/microbiolspec.tbtb2-0004-2015]

28. Orme IM. The mouse as a useful model of tuberculosis. Tuberculosis (Edinb). 2003;83(1-3):112–5. [10.1016/s1472-9792(02)00069-0]

29. Lahey T, Laddy D, Hill K, Schaeffer J, Hogg A, Keeble J, et al. Immunogenicity and Protective Efficacy of the DAR-901 Booster Vaccine in a Murine Model of Tuberculosis. PLoS One. 2016;11(12):e0168521. [10.1371/journal.pone.0168521]

30. Wang P, et al. Immunotherapeutic efficacy of recombinant Mycobacterium smegmatis expressing Ag85B-ESAT6 fusion protein against persistent tuberculosis infection in mice. Hum Vaccin Immunother. 2014. [10.4161/hv.26171]

31. Plumlee CR, Duffy FJ, Gern BH, Delahaye JL, Cohen SB, Stoltzfus CR, et al. Ultra-low Dose Aerosol Infection of Mice with Mycobacterium tuberculosis More Closely Models Human Tuberculosis. Cell Host Microbe. 2021;29(1):68–82.e5. [10.1016/j.chom.2020.10.003]

32. Kaufmann SH. Future vaccination strategies against tuberculosis: thinking outside the box. Immunity. 2010;33(4):567–77. [10.1016/j.immuni.2010.09.015]

33. Andersen P, Woodworth JS. Tuberculosis vaccines--rethinking the current paradigm. Trends Immunol. 2014;35(8):387–95. [10.1016/j.it.2014.04.006]

34. Aagaard C, Hoang T, Dietrich J, Cardona PJ, Izzo A, Dolganov G, et al. A multistage tuberculosis vaccine that confers efficient protection before and after exposure. Nat Med. 2011;17(2):189–94. [10.1038/nm.2285]

35. Nemes E, Geldenhuys H, Rozot V, Rutkowski Kathryn T, Ratangee F, Bilek N, et al. Prevention of M. tuberculosis Infection with H4:IC31 Vaccine or BCG Revaccination. New England Journal of Medicine. 2018;379(2):138–49. [10.1056/NEJMoa1714021]

36. Dockrell HM, Smith SG. What Have We Learnt about BCG Vaccination in the Last 20 Years? Front Immunol. 2017;8:1134. [10.3389/fimmu.2017.01134]

37. Sweeney KA, Dao DN, Goldberg MF, Hsu T, Venkataswamy MM, Henao-Tamayo M, et al. A recombinant Mycobacterium smegmatis induces potent bactericidal immunity against Mycobacterium tuberculosis. Nature Medicine. 2011;17(10):1261–8. [10.1038/nm.2420]

38. Chang Y, Ning H, Liang X, Lu Y, Kang J, Li X, et al. Recombinant Mycobacterium smegmatis expressing antigen 85B for immunotherapy of Mycobacterium tuberculosis-infected mice. Hum Vaccin Immunother. 2025;21(1):2571287. [10.1080/21645515.2025.2571287]

39. Liu W, Xu Y, Yan J, Shen H, Yang E, Wang H. Ag85B synergizes with ESAT-6 to induce efficient and long-term immunity of C57BL/6 mice primed with recombinant Bacille Calmette-Guerin. Exp Ther Med. 2017;13(1):208–14. [10.3892/etm.2016.3944]

40. Cervantes-Villagrana AR, Hernández-Pando R, Biragyn A, Castañeda-Delgado J, Bodogai M, Martínez-Fierro M, et al. Prime-boost BCG vaccination with DNA vaccines based in β-defensin-2 and mycobacterial antigens ESAT6 or Ag85B improve protection in a tuberculosis experimental model. Vaccine. 2013;31(4):676–84. [10.1016/j.vaccine.2012.11.042]

41. Liang Y, Bai X, Zhang J, Song J, Yang Y, Yu Q, et al. Ag85A/ESAT-6 chimeric DNA vaccine induces an adverse response in tuberculosis-infected mice. Mol Med Rep. 2016;14(2):1146–52. [10.3892/mmr.2016.5364]

42. Fan X, Gao Q, Fu R. DNA vaccine encoding ESAT-6 enhances the protective efficacy of BCG against Mycobacterium tuberculosis infection in mice. Scand J Immunol. 2007;66(5):523–8. [10.1111/j.1365-3083.2007.02006.x]

43. Arora SK, Naqvi N, Alam A, Ahmad J, Alsati BS, Sheikh JA, et al. Mycobacterium smegmatis Bacteria Expressing Mycobacterium tuberculosis-Specific Rv1954A Induce Macrophage Activation and Modulate the Immune Response. Front Cell Infect Microbiol. 2020;10:564565. [10.3389/fcimb.2020.564565]

44. Macleod M. Learning lessons from MVA85A, a failed booster vaccine for BCG. Bmj. 2018;360:k66. [10.1136/bmj.k66]

